# Design and Technical Feasibility Testing of a Medical Robot for Flexible Catheter Navigation Inside the Lung Airways

**DOI:** 10.1101/2024.05.01.592024

**Authors:** Lucian Gheorghe Gruionu, Thomas Langø, Håkon Olav Leira, Erlend Fagertun Hofstad, Anca Loredana Udriştoiu, Andreea Valentina Iacob, Cătălin Constantinescu, Cristian Chihaia, Gabriel Gruionu

## Abstract

The integration of medical robots is revolutionizing clinical medicine, especially in procedures requiring precision in instrument manipulation and navigation within the body using medical imaging techniques like fluoroscopy, CT, and MRI. This is particularly challenging in peripheral lung lesion examinations, where guiding long, flexible instruments through the lung airways to the target exposes medical professionals and patients to harmful X-ray radiation. Several robotic approaches exist but there are still shortcomings in terms of their large footprint in the operating room and complex and costly mechanical structure. The goal of our research was to develop and test the early feasibility of RoboCath, an innovative robotic platform designed for the adaptable navigation of long, flexible catheter-like medical instruments through the lung airways. RoboCath seamlessly integrates with existing medical devices, such as bronchoscopes, facilitating access to lower lung regions with precision. Its compact design is ideal for crowded operating rooms, ensures sterilization compatibility, and supports a broad range of procedures, including those requiring intricate instrument manipulation and medical imaging for navigation. This technology significantly reduces the reliance on X-ray, thereby minimizing radiation exposure to both healthcare providers and patients. The RoboCath system represents a significant advancement in the field of medical robotics, offering a novel solution to the challenges of lung lesion biopsy and diagnosis, with potential applications in various open orifice procedures.

## INTRODUCTION

In recent years, robotic systems designed for lung interventions, particularly lung cancer biopsies, have emerged as a focal point of research and development[1–4]. Lung cancer remains one of the leading causes of cancer-related deaths worldwide, underscoring the critical need for early diagnosis and intervention. Traditional methods of lung biopsy, while effective, come with inherent risks and limitations, including the potential for complications and the need for high precision in navigating the complex bronchial pathways. Recent innovations in medical robotics have sought to address these challenges, introducing systems capable of enhancing the accuracy and safety of lung biopsies. Notably, the introduction of a teleoperated robotic bronchoscopy system as described by Chen et al. [5] marks a significant step forward, offering a minimally invasive approach that combines improved precision with spatial flexibility. Similarly, the work of Sganga et al. [6] demonstrates the application of deep learning for real-time localization of a bronchoscope within the lung, facilitating autonomous navigation and enhancing biopsy accuracy.

Further advancements are represented by Kuntz et al. [7] who present the first medical robot capable of autonomously navigating a needle inside living tissue, adeptly maneuvering around obstacles to reach intra-tissue targets. This development not only promises increased safety but also opens the door to new possibilities in lung cancer diagnosis. Additionally, Schreiber et al. [8] designed an open-source 7-axis robotic platform, specifically engineered for CT-guided percutaneous needle biopsy. This system underscores the role of robotic precision and dexterity in performing complex procedures within constrained environments, such as CT scanners.

Despite their advantages, existing clinically available lung robotic solutions occupy a large space in the operating room, are challenging to sterilize, and their manufacturing can be intricate and costly, limiting their accessibility and clinical use. To address these limitations, we aimed to develop a novel solution, the RoboCath with an innovative design that prioritizes a small profile, making it more suitable for crowded operating rooms. It is designed for ease of manufacturing, utilizing 3D printing technology for cost-effective production and rapid iteration of design improvements. Additionally, RoboCath’s compatibility with a wide range of catheter diameters enhances its adaptability for various medical procedures.

## METHODS

The Robocath robot comprises two primary components that make it suitable for use in a sterile environment (Fig. 1). Component 1, the blue case with all the components inside, is reusable and houses two electric motors (3, 4), specifically the Robotis Dynamixel AX 12A, along with the control boards (7), including the OpenCM9.04-C controller and the OpenCM845 expansion board (Fig. 1). This reusable part can be encased in a sterile bag before the procedure. The single-use sterile component incorporates a specialized cartridge containing a pre-mounted 2 mm OD catheter (5), which guarantees its smooth translation movement, and a gear mechanism (9) that ensures the rotation of a 1.2 guide wire (6) passing through the catheter. The medical instrument (5 + 6) used in conjunction with the robot consists of two components: a hollow catheter (5) installed in the cartridge and a guide wire (6) running through the catheter, connected to the gear mechanism (9). When one of the motors (3) rotates the internal component (8) within the cartridge, the instrument undergoes translation, emerging from the robot, passing through the bronchoscope’s working channel, and entering the airways.

**Figure 1.**
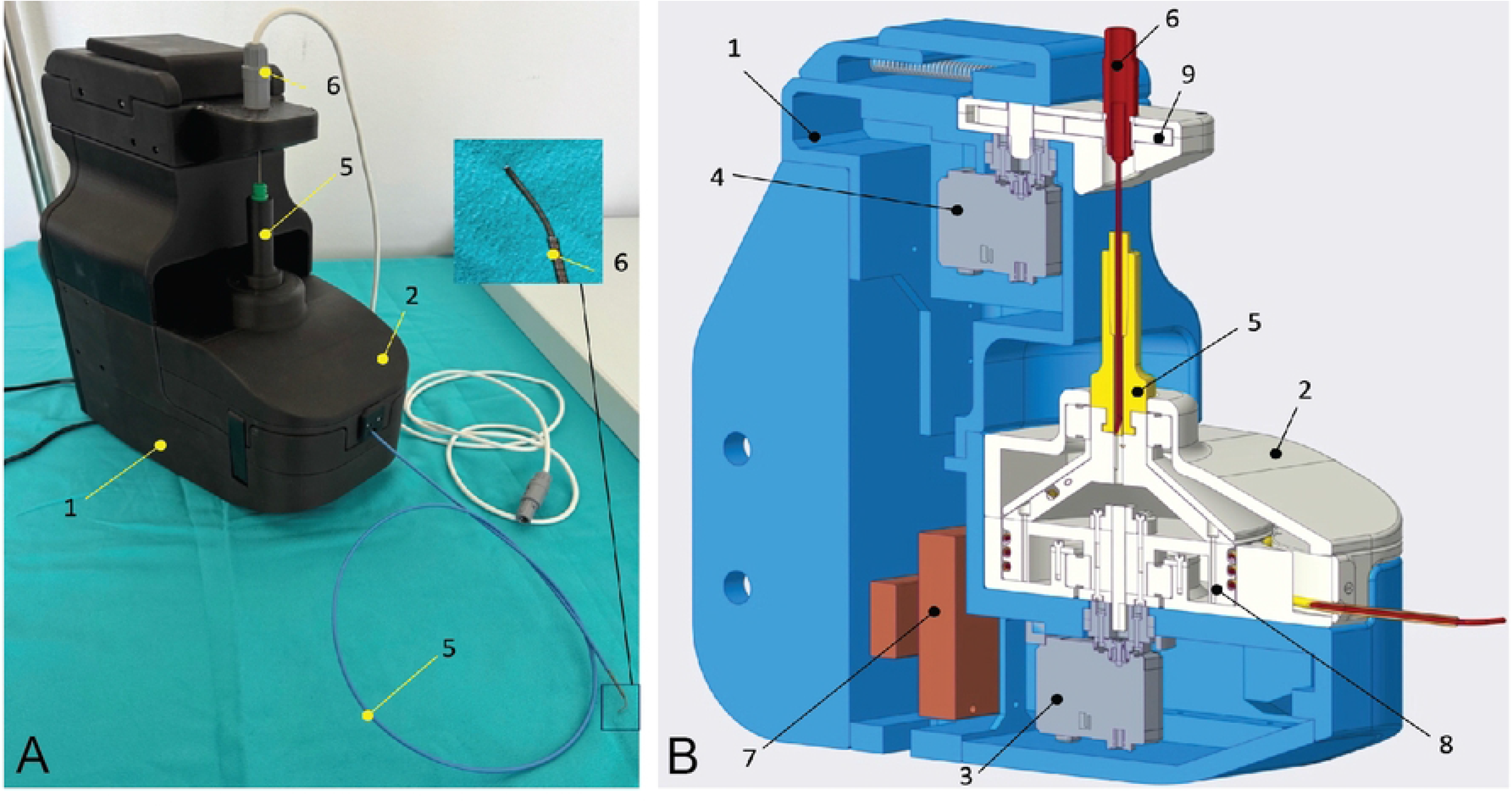
The medical robot for catheter navigation (Robocath): A - 3D printed prototype;1 – reusable component of the 3D printed outer case of the robot; 2-disposable 3D printed component to which medical instruments attach; 3, 4 electronics (not shown in A); 5 – instrument holder; 6 – the handle of a traditional endoscope or a custom-made flexible catheter. B - CAD representation of Robocath; 1 –the reusable outer case which holds the mechanical parts,; 2 – disposable part to which medical instruments attach;3,4 - electric motors; 5,6 - medical instruments attached to the disposable part; 7 - mechanical components;; 8 – gear mechanism for medical instrument’s translation movement; 9-gear mechanism for rotational movement.

By using the second motor (4), Robocath can rotate the proximal part of the guide wire through the gear mechanism. This guide wire has a pre-bent tip at its distal end (detail in the left part of Figure 1), which extends from the working catheter. This tip allows the robot’s user to pinpoint a specific section of the lung airways and navigate close to the target, such as a peripheral nodule. To maintain the rotation angle along the guide wire, it was constructed using a metal coiled spring, as shown in the picture (Fig. 2). The inner channel of this spring houses a 6DOF electromagnetic tracking sensor from Northern Digital, which is employed for navigation purposes.

**Figure 2.**
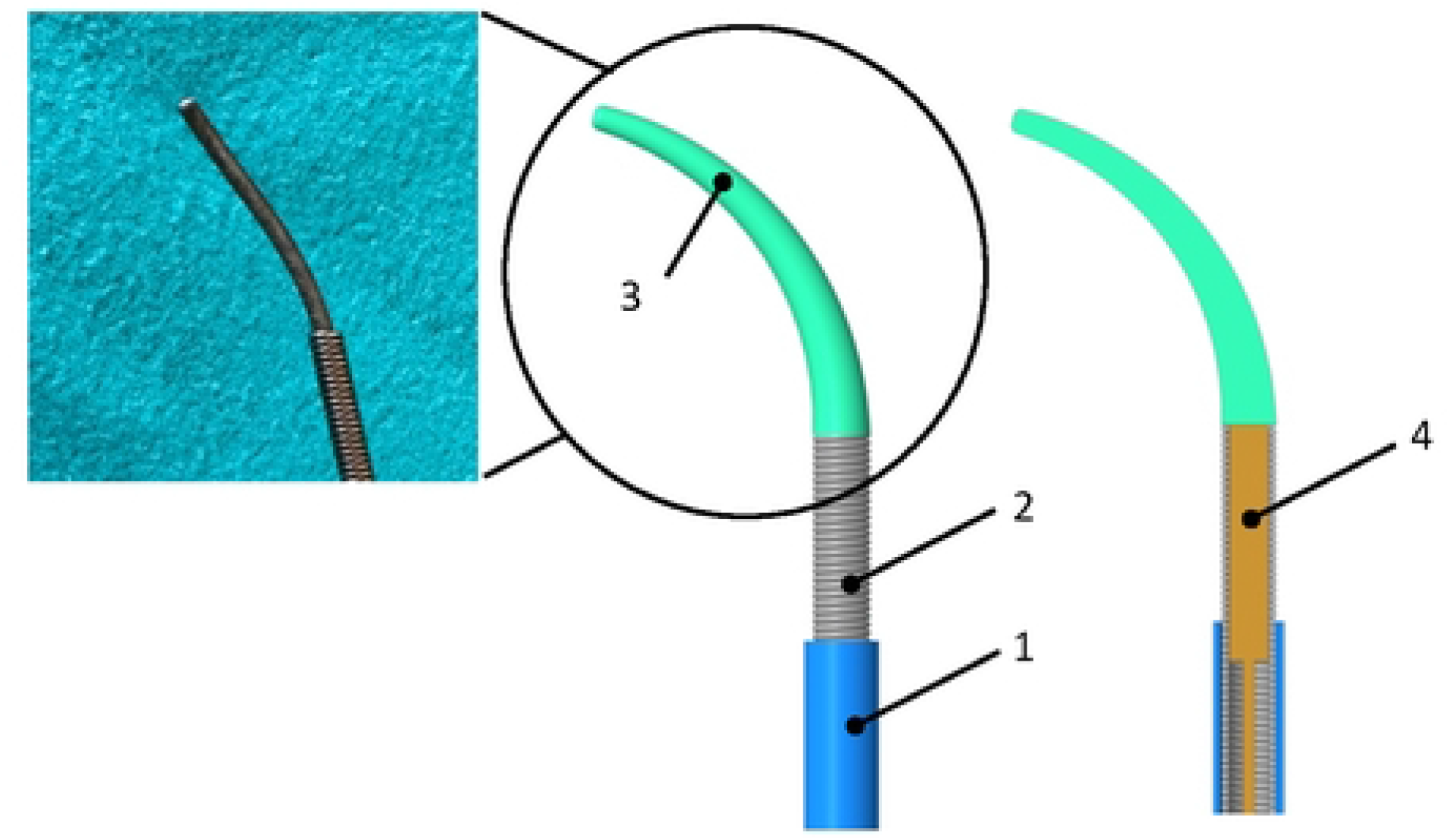
Electromagnetic navigation instrument custom-designed for the Robocath system: 1 – hollow catheter, 2 – metal coiled spring part of the guidewire, 3 – nitinol curved tip, 4 – electromagnetic tracking sensor.

Robocath was designed to be cost-effective and simplified manufacturing through 3D printing. We utilized a Markforged Onyx Pro FDM printer[9], and the components were constructed using micro carbon fiber-filled nylon material. Furthermore, Robocath is used alongside a bronchoscopy system or a custom design catheter. The configuration for a bronchoscopy procedure employing the Robocath system is depicted in Figure 3.

**Fig. 3.**
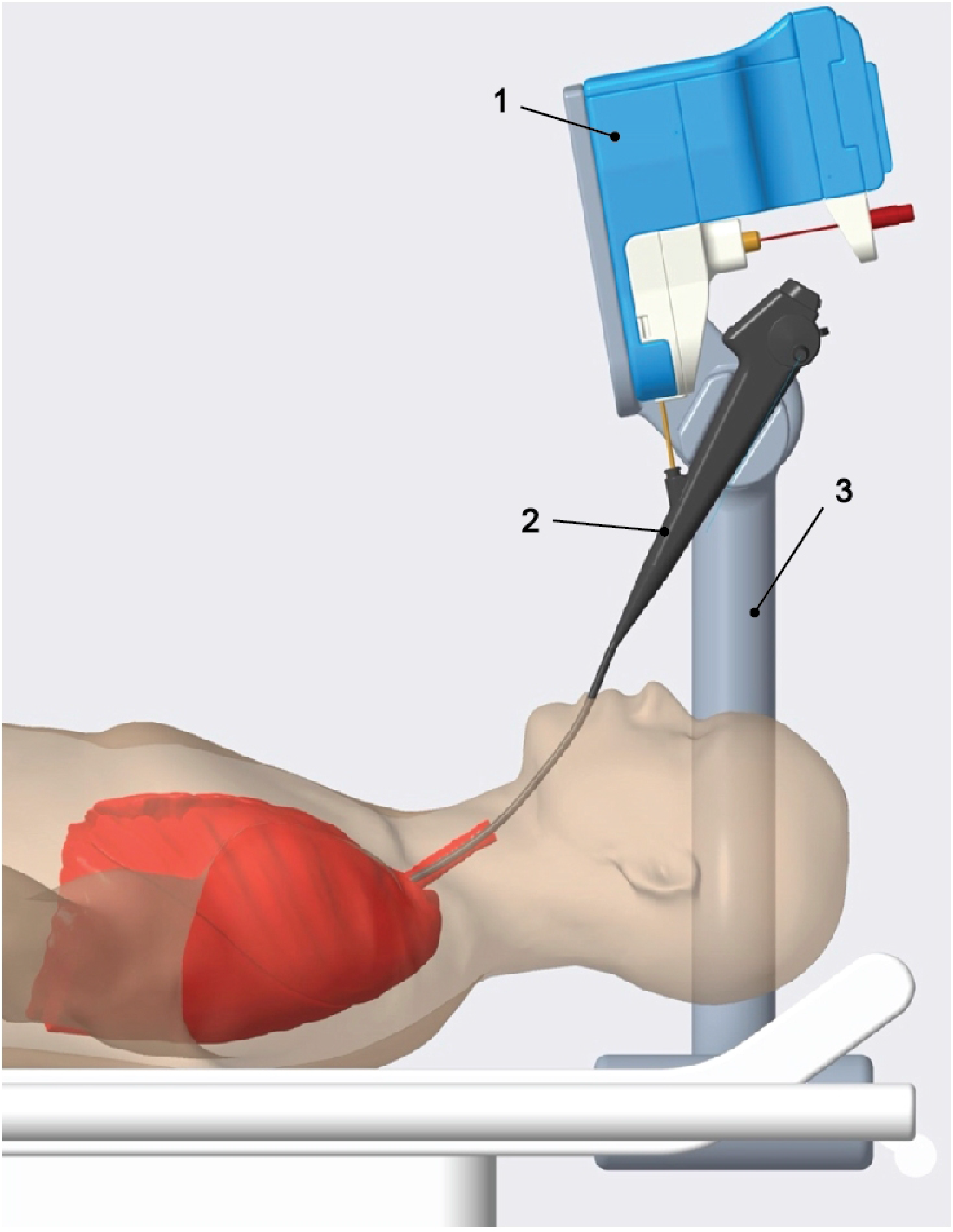
The placement of the Robocath device during the bronchoscopy procedure: 1-the robot is placed over and away from the patient’s head for easy access to the patient’s airways, 2 – the bronchoscope or a flexible catheter is precisely inserted and rotate by the robot, 3 – an adjustable arm allows repositioning of the Robocath device according to the patient’s position.

### The procedure involves the following steps

- Initially, the physician uses the patient’s CT scans and Fraxinus software to locate a lesion on the lung’s periphery that requires further investigation and a biopsy. This software automatically identifies both the airways and the suspected lesion to be investigated. The physician confirms the software’s findings and utilizes the same software to plan the biopsy procedure and compute the optimal route from the trachea to the target lesion.
- The physician then conducts a standard bronchoscopy procedure using a bronchoscope, advancing towards the target until the bronchial diameter becomes too small for further advancement of the bronchoscope.
- A nurse brings the adjustable handle equipped with the robot near the bronchoscope by sliding it along the rail on the procedure table and adjusting the handle’s height and rotation. This positioning allows the bronchoscope handle to be secured without moving it, as illustrated in Figure 2.
- The single-use components, including the medical instrument, catheter, and guidewire of the Robocath system (as shown in Figures 1, 2, 5, and 6), are removed from their sterile packaging and installed in the robot. The robot’s design permits its placement and operation within a sterile bag during the procedure.
- After mounting the bronchoscope onto the handle and inserting the catheter with the guidewire into the bronchoscope’s working channel and connecting the AURORA tracking system, the physician can navigate the catheter using the Robocath robot through its software interface (Fig. 4).
- The Fraxinus software displays the position of the guidewire tip in relation to a 3D model of lung anatomy, utilizing the AURORA system and the electromagnetic sensor [10] to provide real-time location and orientation.
- The physician uses a specialized user interface to control the robot’s motors, allowing for the catheter to be moved and the guidewire with a curved, flexible tip, to be rotated. This enables the selection of a specific bronchial path that was calculated during the planning phase.
- Upon positioning the catheter near the target, the physician withdraws the guidewire and inserts a biopsy needle through the catheter to perform the biopsy, guided by the robot.

**Fig. 4.**
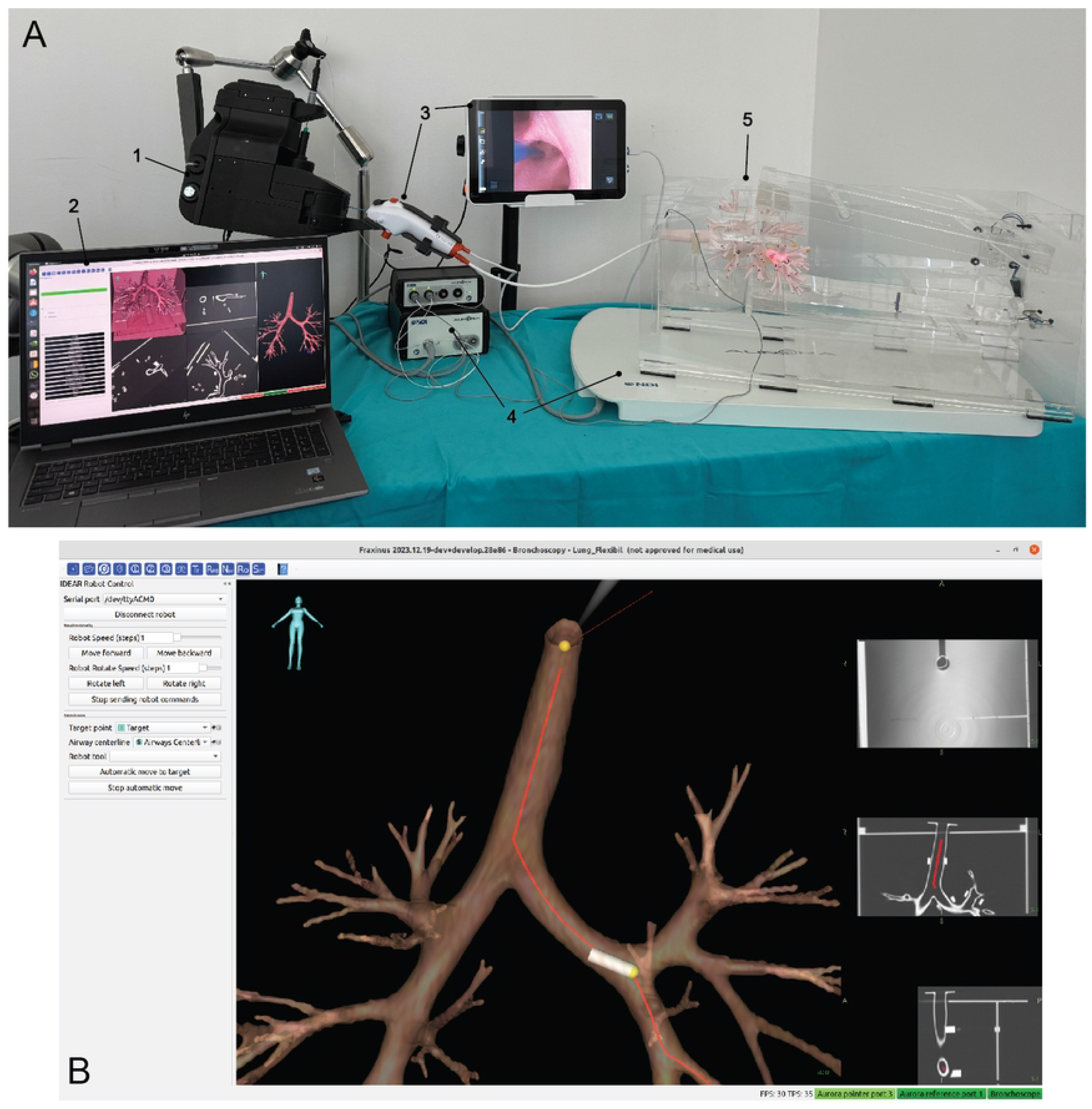
The Robocath system testing set-up. A: The workstation displays the Fraxinus navigation software platform with the 3-D model of the lung phantom. The AMBU Scope Broncho bronchoscope is control through a table interface. The NDI Aurora electromagnetic navigation system is located under the phantom and used to trace the catheter in the lung airways. B: A screen shot of theFraxinus interface for the robotic procedure.

**Figure 5.**
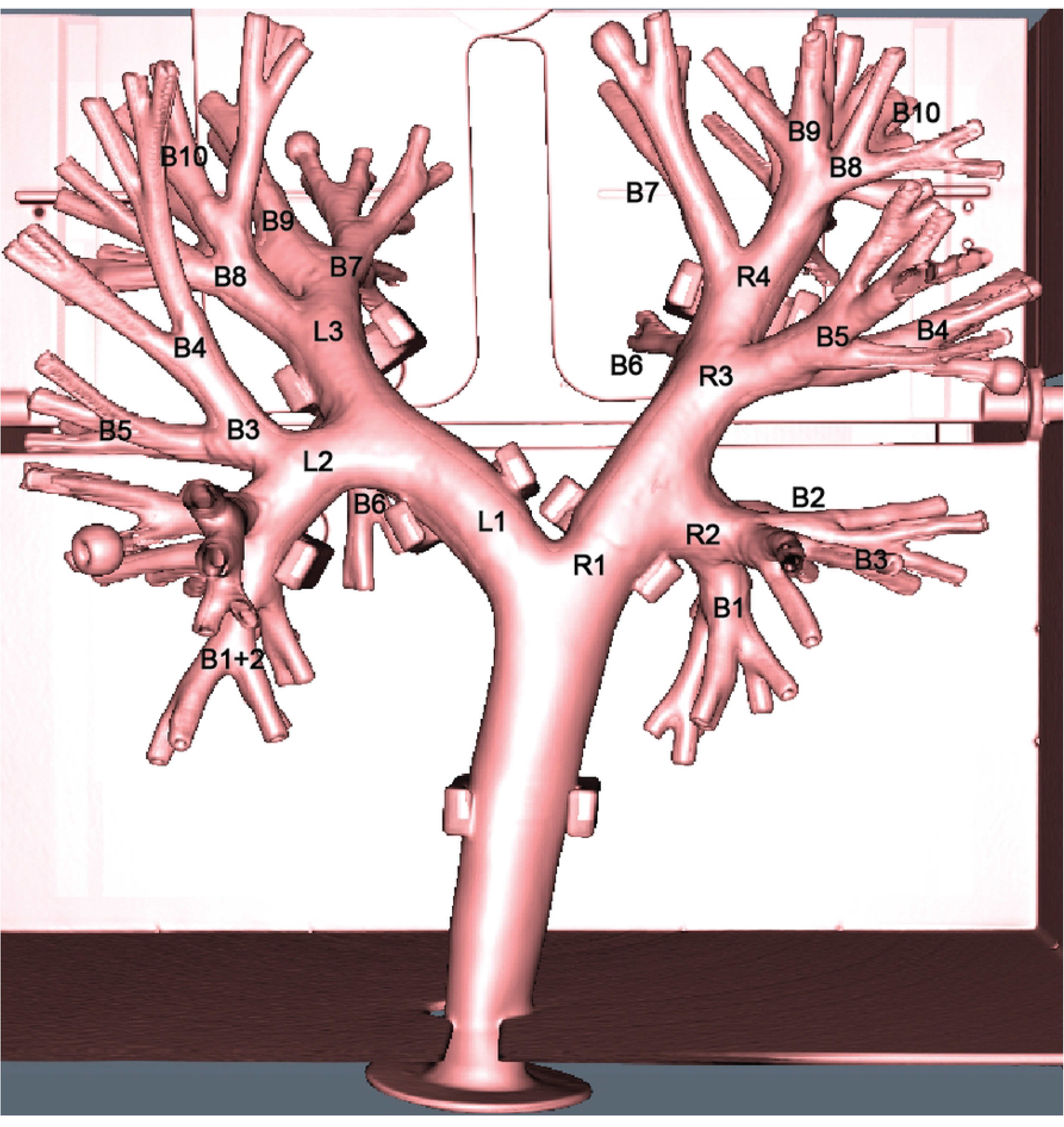
The 3D rendering of the airways phantom with clinically relevant airway labels (see Table 1).

**Table 1.**
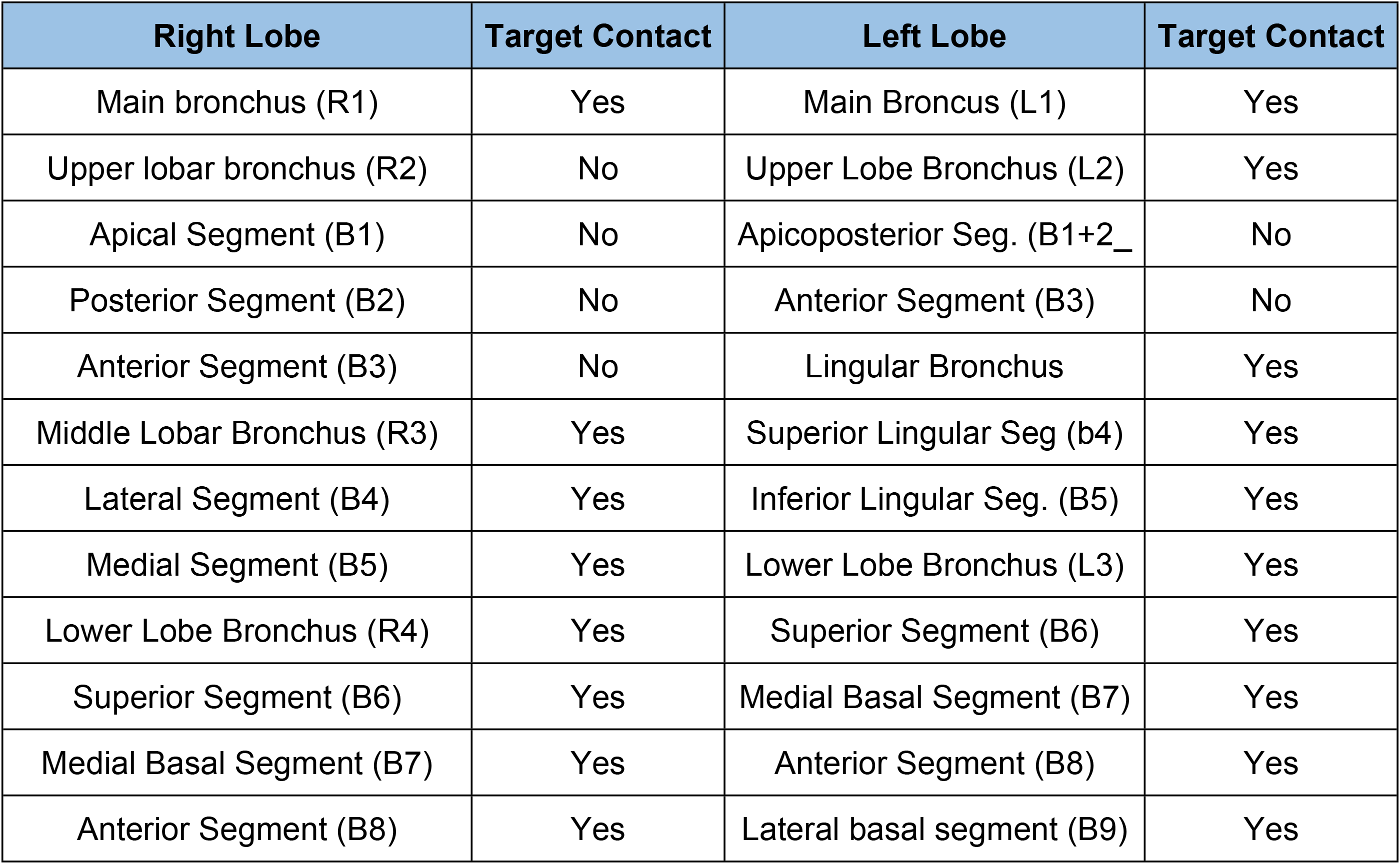
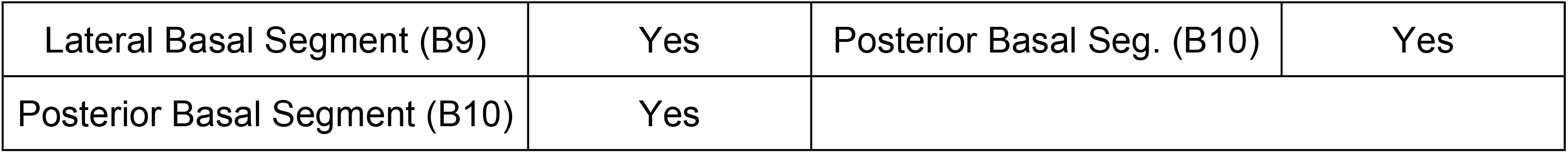
Target contact for lung bronchial tree branches using the Robocath system during feasibility training.

## RESULTS

The primary goal of the initial testing of the Robocath was to assess its capability to navigate the guidewire to all bronchial locations within a realistic and widely accepted bronchoscopy phantom, the KOKEN Bronchoscopy Simulator[11]. The phantom was first scanned by computed tomography (CT) similar to clinical imaging procedure. The 2-D CT scan series were uploaded into the publicly available Fraxinus software platform developed by our team to generate a 3-D map of the lung anatomy model and allow computer and electromagnetic guidance[12].

In the CT scan series, a hypothetical target, designated as a “lesion,” was identified at the termination of each model airway for procedural planning purposes. The software was tasked with determining the most efficient route from the trachea, serving as the entry point, to each “lesion.” To synchronize the physical model with its digital reconstruction derived from the CT scans, a single external marker was placed close to the trachea’s entrance. For the experimental trials, the AmbuaScope 4 Broncho bronchoscopy system was employed, selecting the largest variant within the series that features a 2.6 mm channel width for instruments. The configuration of the laboratory bench designated for these tests is showcased in Figure 4.

The procedure began with the bronchoscope being manually navigated towards the “lesion” until the point where the airways narrowed significantly, usually at the third or fourth bifurcation. At this juncture, the bronchoscope was positioned on a stand that also accommodated the Robocath. The catheter, protruding from its cartridge, was then fed through the working channel of the bronchoscope. From this moment, the operator managed the Robocath through an intuitive interface to advance the catheter beyond the end of the bronchoscope, employing the Fraxinus software for directional guidance to the “lesion.” Upon arriving at the designated target, the guide wire equipped with the EM sensor was retracted, making room for the insertion of a biopsy instrument into the catheter for sample collection.

This approach enabled users to effectively access numerous targets within the peripheral branches of the lung, manipulating the instruments’ movements, translations, and rotations via the Robocath integrated with the bronchoscope (Table 1).

## CONCLUSIONS

The primary clinical application of the Robocath system is its ability to position a working catheter or bronchoscope in close proximity to a target lesion inside the lung airways. This catheter can be utilized to image or introduce a biopsy tool safely in the proximity of the lesion. Users can navigate the catheter with the assistance of the robot, imaging software, and electromagnetic tracking software, without the need for fluoroscopy scans, thus mitigating the occupational hazards associated with interventional radiology. Due to its coiled spring design, the guidewire within the catheter maintains the rotation angle consistently from the proximal to the distal end, facilitating path-following in a tubular system.

Robocath was engineered with a focus on cost-effectiveness and streamlined manufacturing, employing 3D printing technology to accomplish these objectives. The utilization of 3D printing allows for optimized production processes, minimized material waste, and enhanced supply chain efficiency. Consequently, this approach contributes to a reduction in the overall cost of the device. By offering a more economically viable robotic-assisted platform for coronary interventions, Robocath aims to facilitate wider adoption in clinical settings and broaden patient access to advanced medical technologies.

RoboCath distinguishes itself from existing technologies through its innovative design that addresses key operational and practical challenges in lung interventions. Its small profile makes it especially suitable for the crowded environment of operating rooms, where space is at a premium. Unlike current clinically approved medical robots, RoboCath has been engineered for ease of manufacture, primarily utilizing 3D printing technology with micro carbon fiber-filled nylon material, which not only simplifies production but also reduces costs. Moreover, RoboCath’s adaptability to a wide range of catheter diameters sets it apart, offering unparalleled versatility for different diagnostic and therapeutic needs within the pulmonary space.

Additional tests are currently underway to assess the positional precision and procedural time using multiple catheters, comparing the Robocath system with a traditional intervention that relies on fluoroscopy for guidance. Future studies will focus on further optimization and pre-clinical testing.

## AKNOWLEDGMENTS

The research leading to these results has received funding from Norwegian Financial Mechanism 2014-2021 under the project RO-NO-2019-0138, 19/2020 “Improving Cancer Diagnostics in Flexible Endoscopy using Artificial Intelligence and Medical Robotics” IDEAR, Contract No. 19/2020.

## Notes

### Competing Interest Statement

The authors have declared no competing interest.

## REFERENCES

1. Chaddha U, Kovacs SP, Manley C, Hogarth DK, Cumbo-Nacheli G, Bhavani SV, et al. Robot-assisted bronchoscopy for pulmonary lesion diagnosis: results from the initial multicenter experience. BMC Pulm Med. 2019 Dec 11;19:243.

2. Diddams MJ, Lee HJ. Robotic Bronchoscopy: Review of Three Systems. Life (Basel). 2023 Jan 28;13(2):354.

3. Zhang C, Xie F, Li R, Cui N, Herth FJF, Sun J. Robotic‐assisted bronchoscopy for the diagnosis of peripheral pulmonary lesions: A systematic review and meta‐analysis. Thorac Cancer. 2024 Jan 29;15(7):505–12.

4. Hammad Altaq H, Parmar M, Syed Hussain T, Salim DJ, Chaudry FA. The Use of Robotic-Assisted Bronchoscopy in the Diagnostic Evaluation of Peripheral Pulmonary Lesions: A Paradigm Shift. Diagnostics (Basel). 2023 Mar 9;13(6):1049.

5. Chen XY, Xiong X, Wang X, Li P, Wang S, Akinyemi T, et al. Design of a Teleoperated Robotic Bronchoscopy System for Peripheral Pulmonary Lesion Biopsy.

6. Sganga J, Camarillo DB. Autonomous Driving in the Lung using Deep Learning for Localization.

7. Kuntz A, Emerson M, Ertop TE, Fried I, Fu M, Hoelscher J, et al. Autonomous Medical Needle Steering In Vivo [Internet]. arXiv; 2022 [cited 2024 Mar 22]. Available from: http://arxiv.org/abs/2211.02597

8. Schreiber DA, Shak DB, Norbash AM, Yip MC. An Open-Source 7-Axis, Robotic Platform to Enable Dexterous Procedures within CT Scanners. In: 2019 IEEE/RSJ International Conference on Intelligent Robots and Systems (IROS) [Internet]. 2019 [cited 2024 Mar 24]. p. 386–93. Available from: http://arxiv.org/abs/1903.04646

9. Onyx Pro Professional 3D Printer | Markforged [Internet]. [cited 2024 Mar 24]. Available from: https://markforged.com/3d-printers/onyx-pro

10. Aurora - NDI’s premier electromagnetic tracking solution [Internet]. NDI. [cited 2024 Mar 24]. Available from: https://www.ndigital.com/electromagnetic-tracking-technology/aurora/

11. Bronchoscopy Training Model LM-092 | Koken Co., Ltd. [Internet]. [cited 2024 Mar 24]. Available from: https://www.kokenmpc.co.jp/english/products/educational_medical_models/anatomical/lm-092.html

12. SINTEFMedtek/Fraxinus [Internet]. SINTEF Medical Technology; 2023 [cited 2024 Mar 24]. Available from: https://github.com/SINTEFMedtek/Fraxinus

